# Contribution to the genus Sympetrum Newman, 1833 (Odonata, Anisoptera) from extreme SW Romania

**DOI:** 10.1101/2022.10.31.514645

**Authors:** Anda Felicia Babalean

## Abstract

The paper presents new data on species composition and distribution of the genus Sympetrum in the extreme SW part of Romania. Two species, *Sympetrum fonscolombii* (Selys, 1840) and *Sympetrum striolatum* (Charpentier, 1840) are added to those of the literature - *Sympetrum sanguineum* (Müller, 1764) and *Sympetrum meridionale* (Selys, 1841) (Cîrdei & Bulimar 1965). The sampled area was enlarged with new collecting sites.

A brief description of species is given, especially the distinctive characters, including the hamular processes and the vulvar scale. The bifid aspect of the vulvar scale in *S. meridionale* indicates that the genus Sympetrum has a problem at least of species diagnosability.

## Introduction

Extreme SW Romania refers to Dolj county. The geographical morphology of this region is diverse and comprises low and high plains, also the hilly units of the Getic Piedmont. The hydrographic network is also rich and diverse, consisting of surface waters (rivers, lakes) and an important subterranean aquifer between which there are reciprocal relations (Pleniceanu 1999, Savin 2000).

The Sympetrum fauna of this region is understudied in terms of species account and distribution, especially since in the last 30 years the area has been subjected to anthropic stress. Cîrdei & Bulimar (1965) give only 2 species – *Sympetrum sanguineum* (Müller, 1764) and *Sympetrum meridionale* (Selys, 1841) from 2 localities – Căciulătești and Sadova.

The aim of this paper is to enlarge the knowledge on Sympetrum species in this region.

## Material and method

Sympetrum species were collected from sites in:

Băilești Plain

- Balta Cilieni (Cilieni Lake/Pool), near Băilești locality
- Maglavit Lake, near Maglavit locality

Romanați Plain

- Lișteava Pool, near Căciulătești locality
- several pools and lakes in Craiova locality: Romanescu Park, Balta Craioviței (Craiovița Pool), Lacul Tanchistului (Tankman Lake/Pool)

Getic Piedmont

- Filiași Central Lake

The specimens were collected between 16 June and 18 October 2022 with the entomologic net, in 80° to absolute ethanol.

## Results

Systematic account

Ord. Odonata

Subord. Anisoptera

Fam. Libellulidae

1) *Sympetrum fonscolombii* (Selys, 1840) – Figs. 1 – 3

**Fig. 1.**
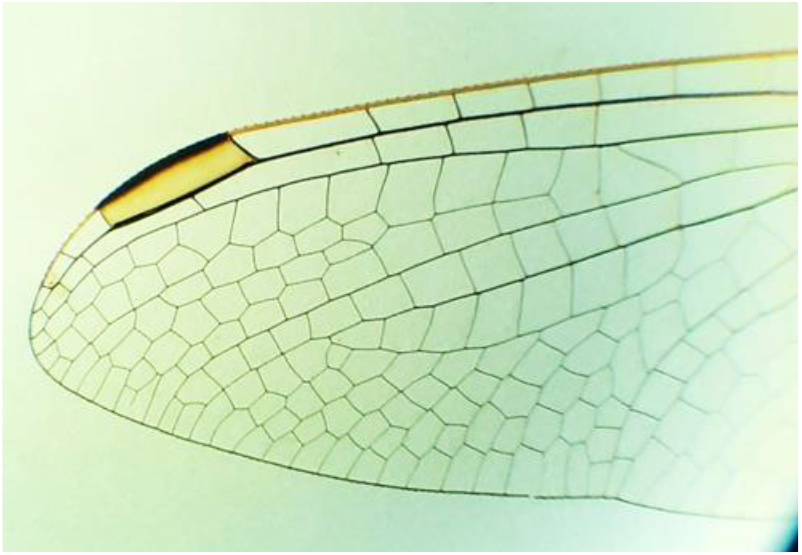
*Sympetrum fonscolombii* – pterostigma (specimen from Maglavit, 16 June 2022)

Lișteava Pool – 28 July, 1♀

Maglavit Lake – 16 June, 1♂

Brief description: striped legs in black and yellow; wings with large amber patches, pterostigma yellow with black borders (Fig. 1); hamular processes (Fig. 2): inner anterior process short, outer posterior process wide; vulvar scale (Fig. 3) with two lobes, widely separated, the S9 sternite has two digitiform prominences.

**Fig. 2.**
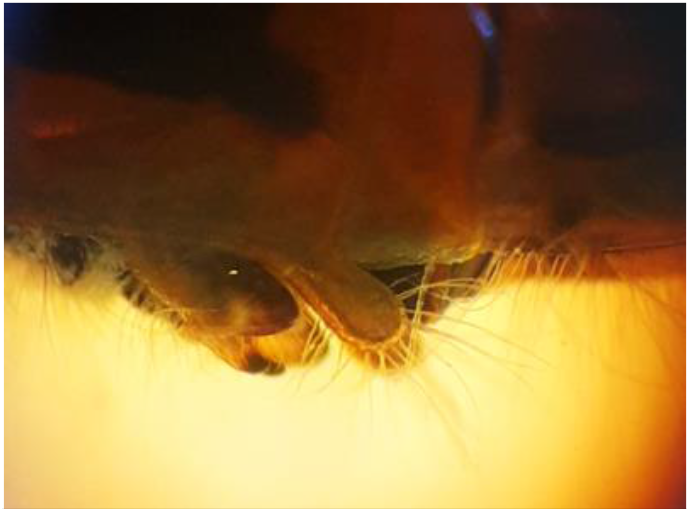
*Sympetrum fonscolombii* – the hamular processes (specimen from Maglavit, 16 June 2022)

**Fig. 3.**
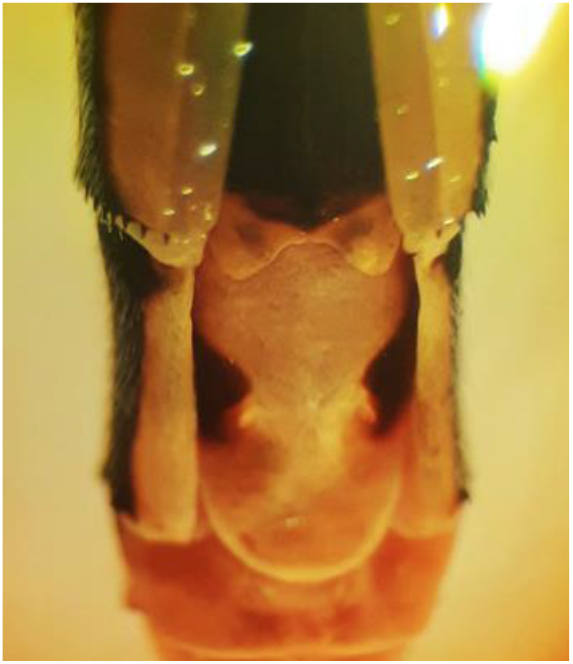
*Sympetrum fonscolombii* – the vulvar scale (specimen from Lișteava, 28 July 2022)

2) *Sympetrum striolatum* (Charpentier, 1840) – Figs. 4 – 6

**Fig. 4.**
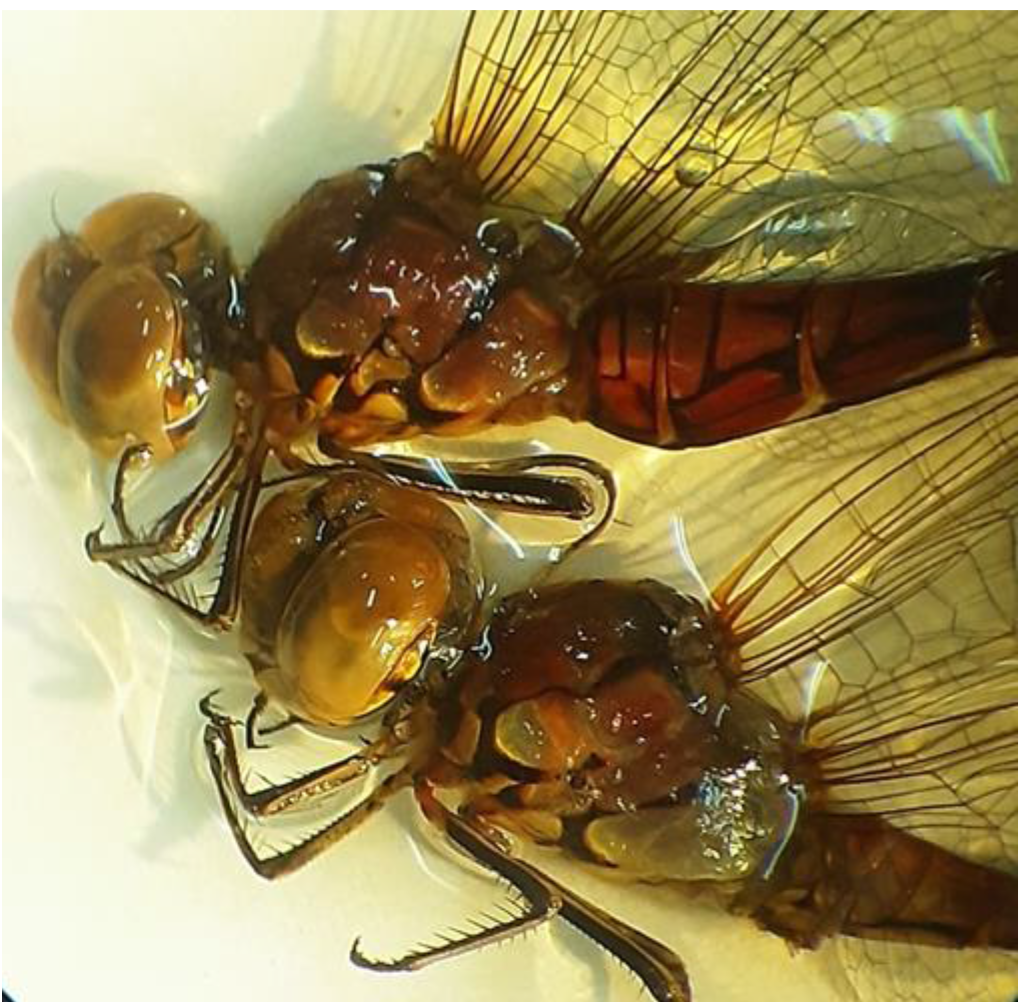
*Sympetrum striolatum* – male and female general habitus, specimens from Craiovița Pool, 25 September 2022

Craiova: Romanescu Park – 22 September, 1♂; Craiovița Pool, 25 September, 3♂♂, 1♀;

Tankman Pool, 16 October, 8♂♂, 9♀♀

Filiași Central Lake – 7 October, 2♂♂

Cilieni Pool Băilești – 18 October, 1 ♂

Brief description: striped legs in black and yellow; lateral synthorax with two large greenish-yellow bands (Fig. 4); hamular processes (Fig. 5) nearly equal: the inner anterior process slender, the outer posterior process wide; vulvar scale (Fig. 6) moderately prominent, with a wide undulation, the sternite below with two digitiform prominences.

**Fig. 5.**
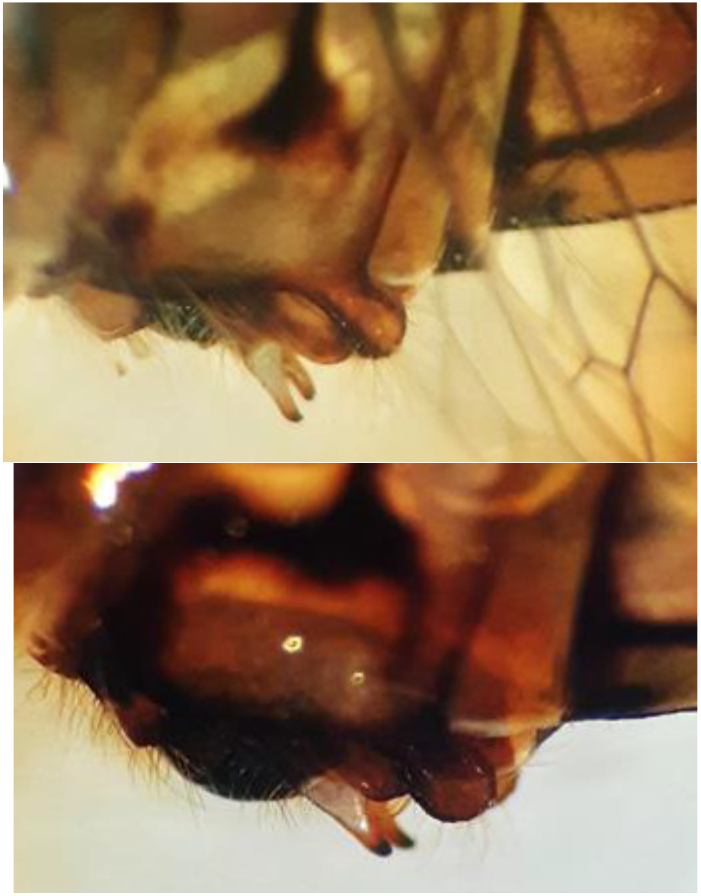
*Sympetrum striolatum* – the hamular processes, specimens from Craiovița Pool, 25 September 2022 (up) and Tankman Lake, 16 October 2022 (down)

**Fig. 6.**
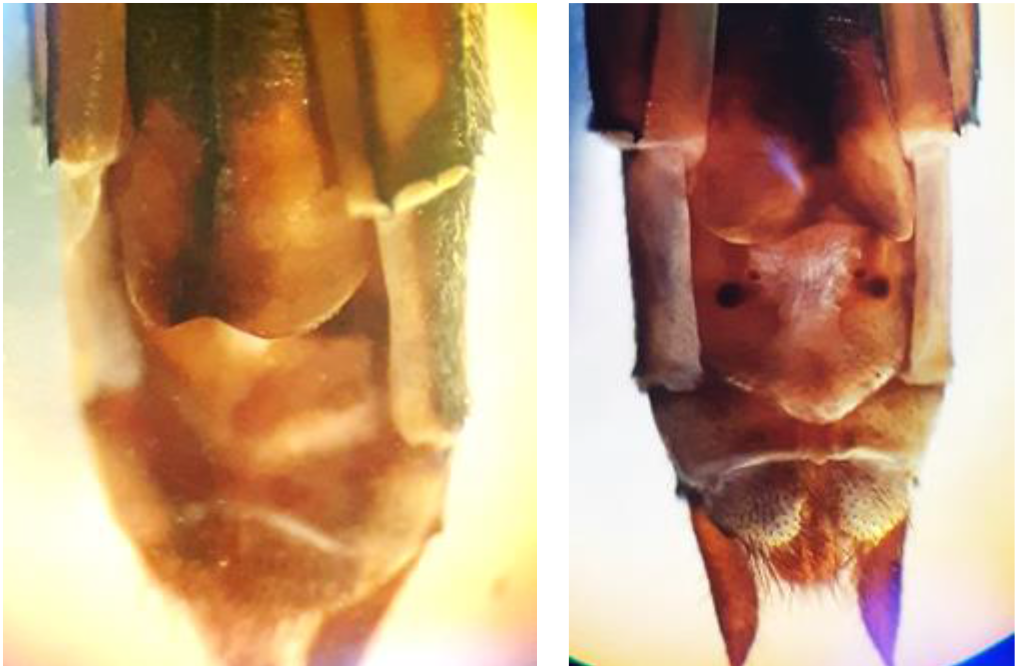
*Sympetrum striolatum* – vulvar scale, specimens from Craiovița Pool, 25 September 2022 (left) and Tankman Lake, 16 October 2022 (right)

3) *Sympetrum meridionale* (Selys, 1841) – Figs. 7 – 11

**Fig. 7.**
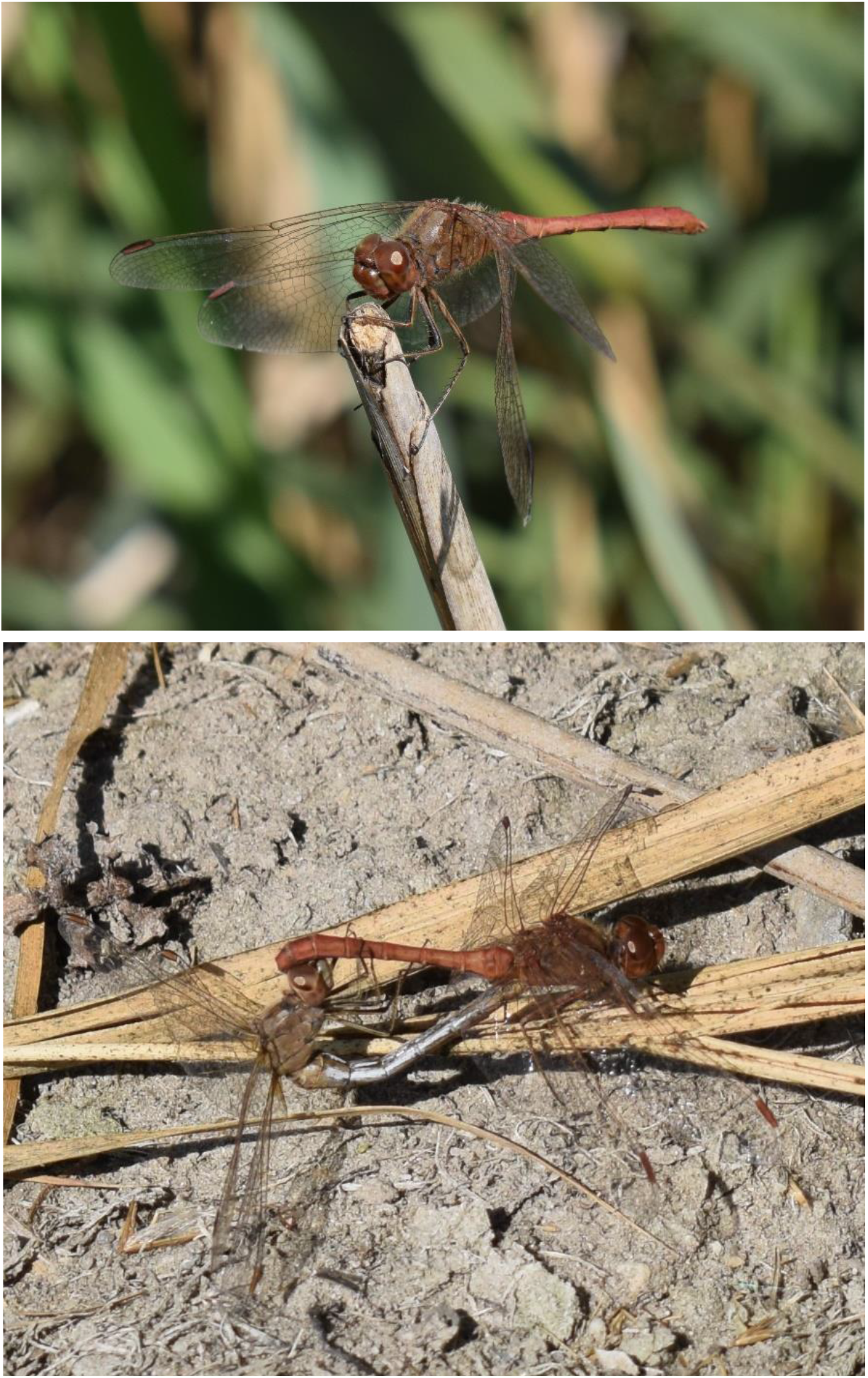
*Sympetrum meridionale* in living, specimens from Cilieni Pool Băilești, male and couple

Cilieni Pool, Băilești – 18 October, 3♂♂, 2♀♀

Brief description

Male (Figs. 7, 8, 9): legs mainly yellow (outer surfaces of femora and tibiae yellow, inner surfaces black); head: frons reddish brown in living (Fig. 7) and yellow-orange in alcohol and under the light of binocular, clypeus area fragmented, labrum reddish, mandibula greenish in the upper part and reddish in the lower part, labium red (Figs. 8, 9); wings: the amber patch small, pterostigma brown; thorax: reddish brown on dorsal and lateral, the lateral black sutures unmarked / slightly marked, the pattern of the yellow spots on lateral thorax is distinctive, the mesepimeron with a conspicuous mark in the shape of the letter “delta” (black core, yellow margins) near the junction of mid coxa (Figs. 8, 9), yellow spiracle with black border, a black point above the spiracle; the abdomen reddish-brown with S1-S2 and S8-S10 deeper red, S8-S10 obviously more dilated (vespoid appearance), S8-S9 without dorsal black spots (Figs. 8, 9); hamular processes unequal, the outer posterior process long triangular shape and shorter than the inner anterior process which ends in a black hooked tip (Figs. 8, 9).

**Fig. 8.**
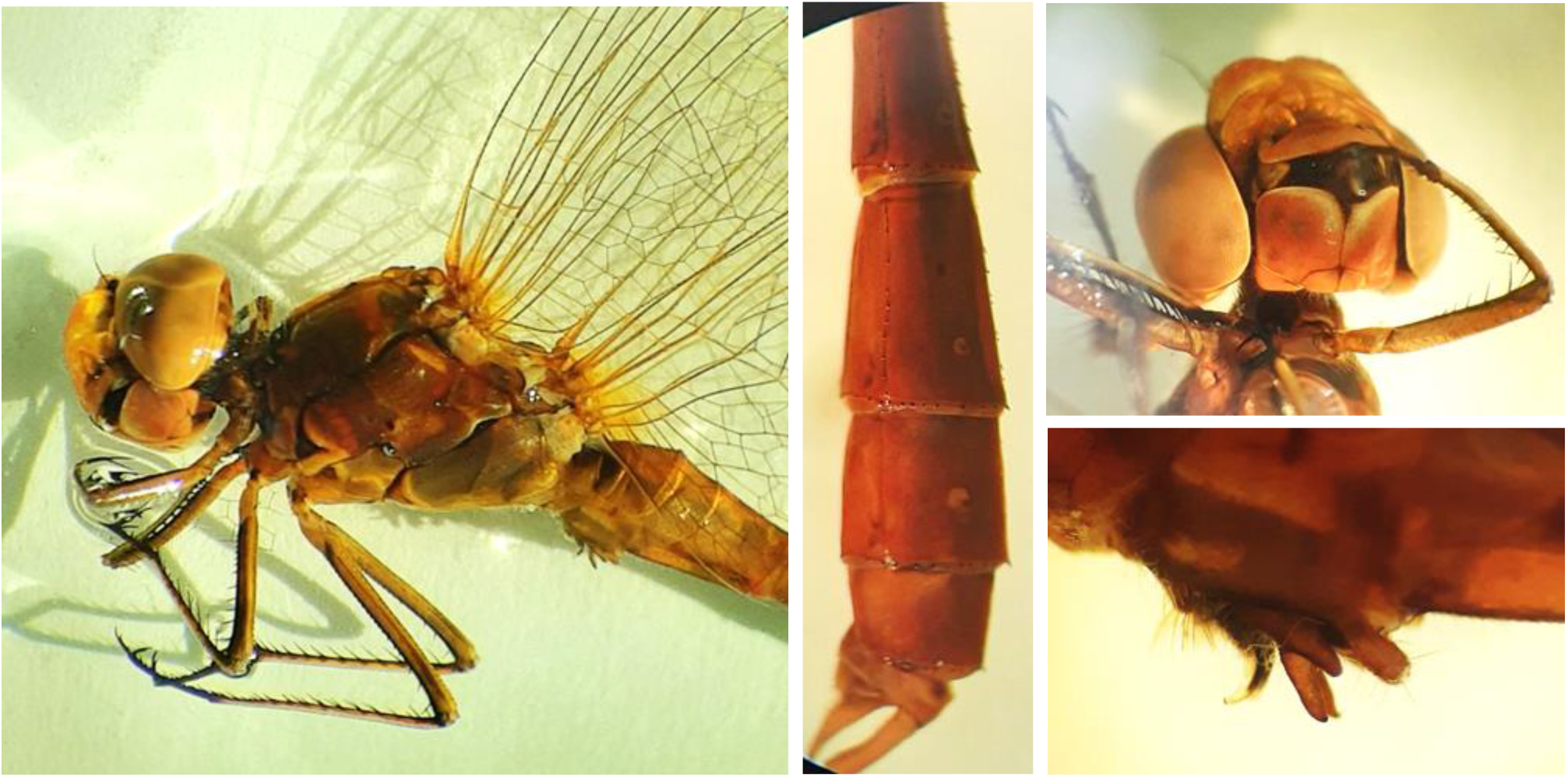
*Sympetrum meridionale*, male specimen from Cilieni Pool Băilești General habitus, the last segments of abdomen, face and hamular processes

**Fig. 9.**
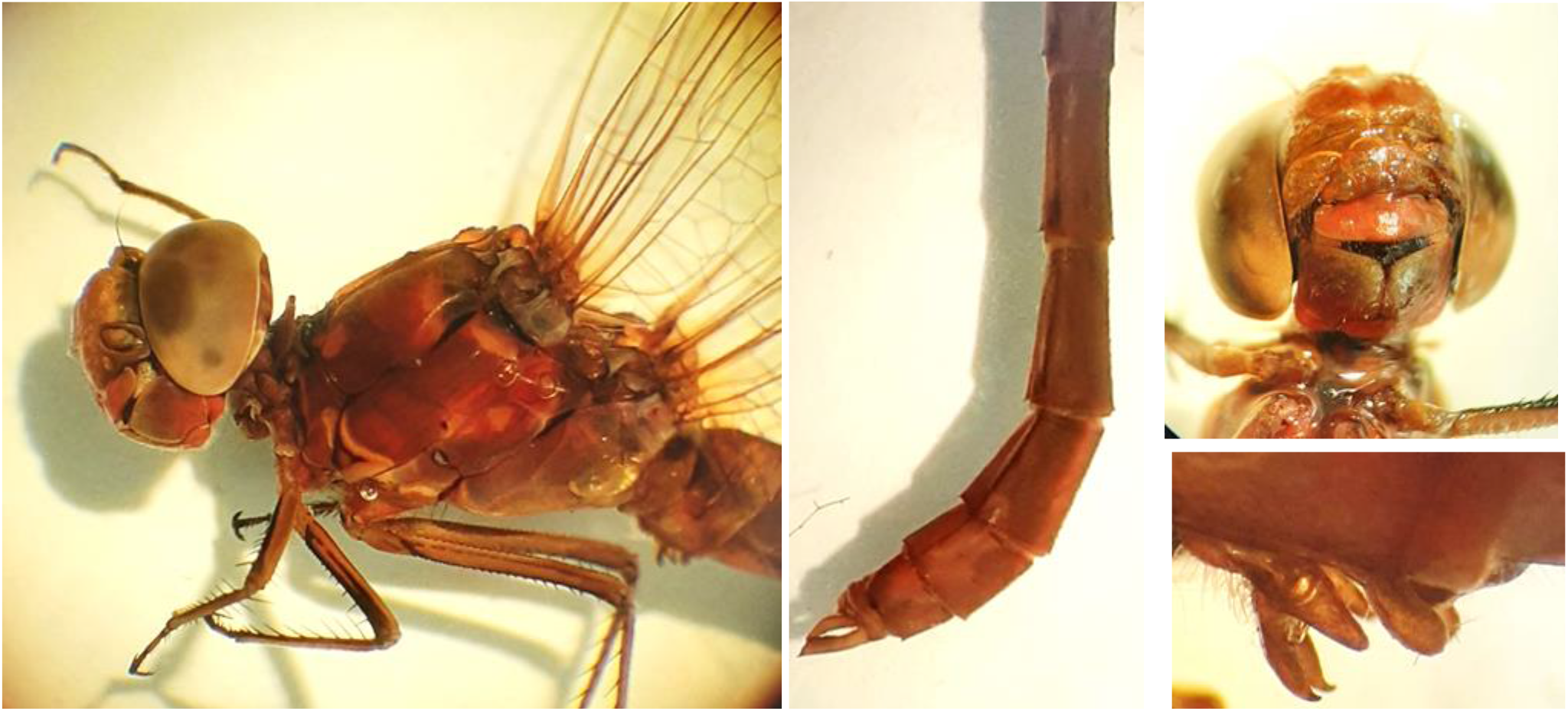
*Sympetrum meridionale*, another male specimen from Cilieni Pool Băilești General habitus, the last segments of abdomen, face and hamular processes

Female (Figs. 7, 10, 11): paler in colour than male, with a greyish tint; legs as in males; head: face yellow greenish without the reddish coloration characteristic of males, except the frons and postclypeus which show some red; thorax with the delta mark present; abdomen without inflated areas, lighter in colour than males, blue to metallic pruinescent in living (Fig. 7); the vulvar scale visible in lateral view, bifid in ventral view (Figs. 10, 11).

**Fig. 10.**
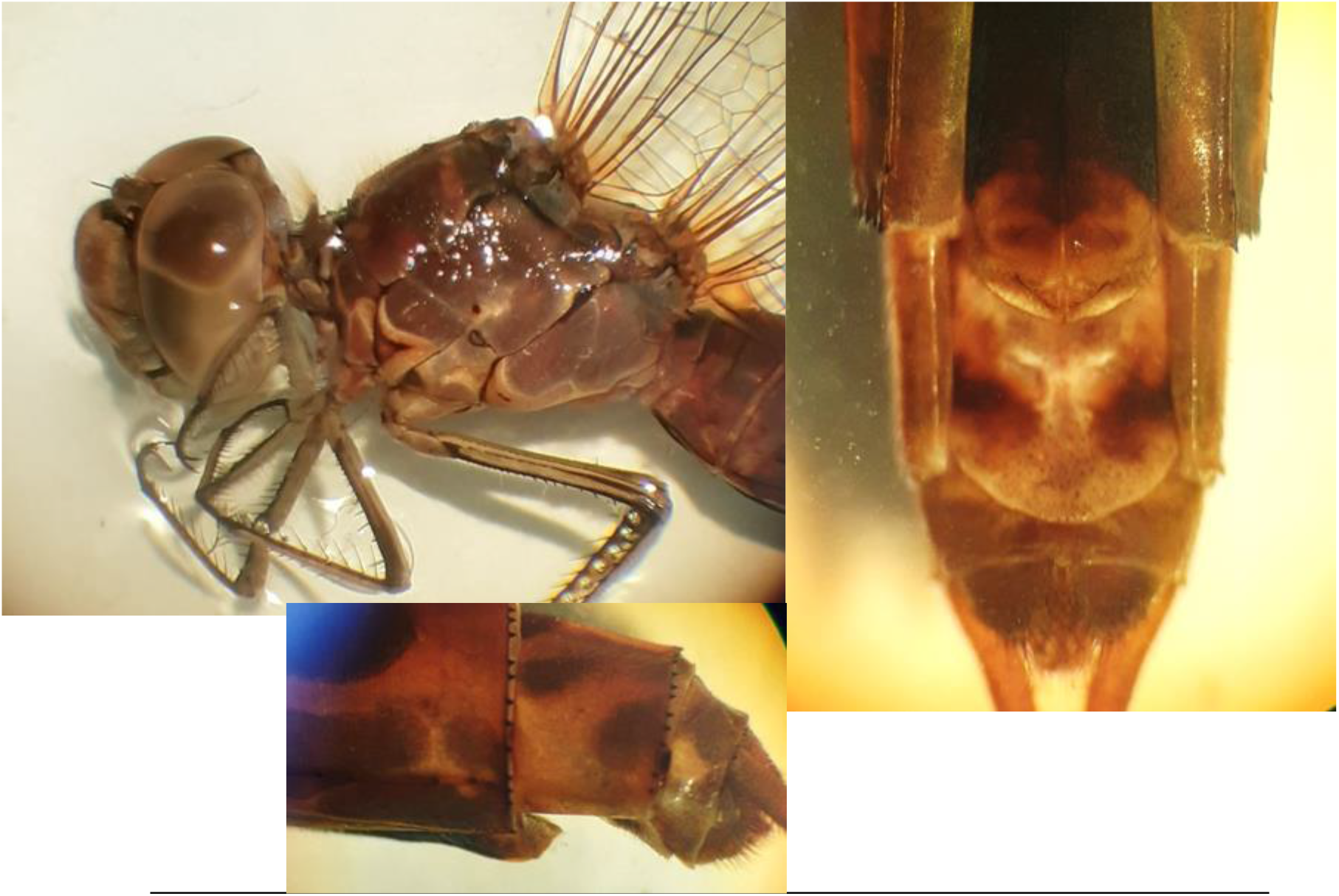
*Sympetrum meridionale* female, the first specimen from Cilieni Pool Băilești General habitus, vulvar scales in lateral and ventral view

**Fig. 11.**
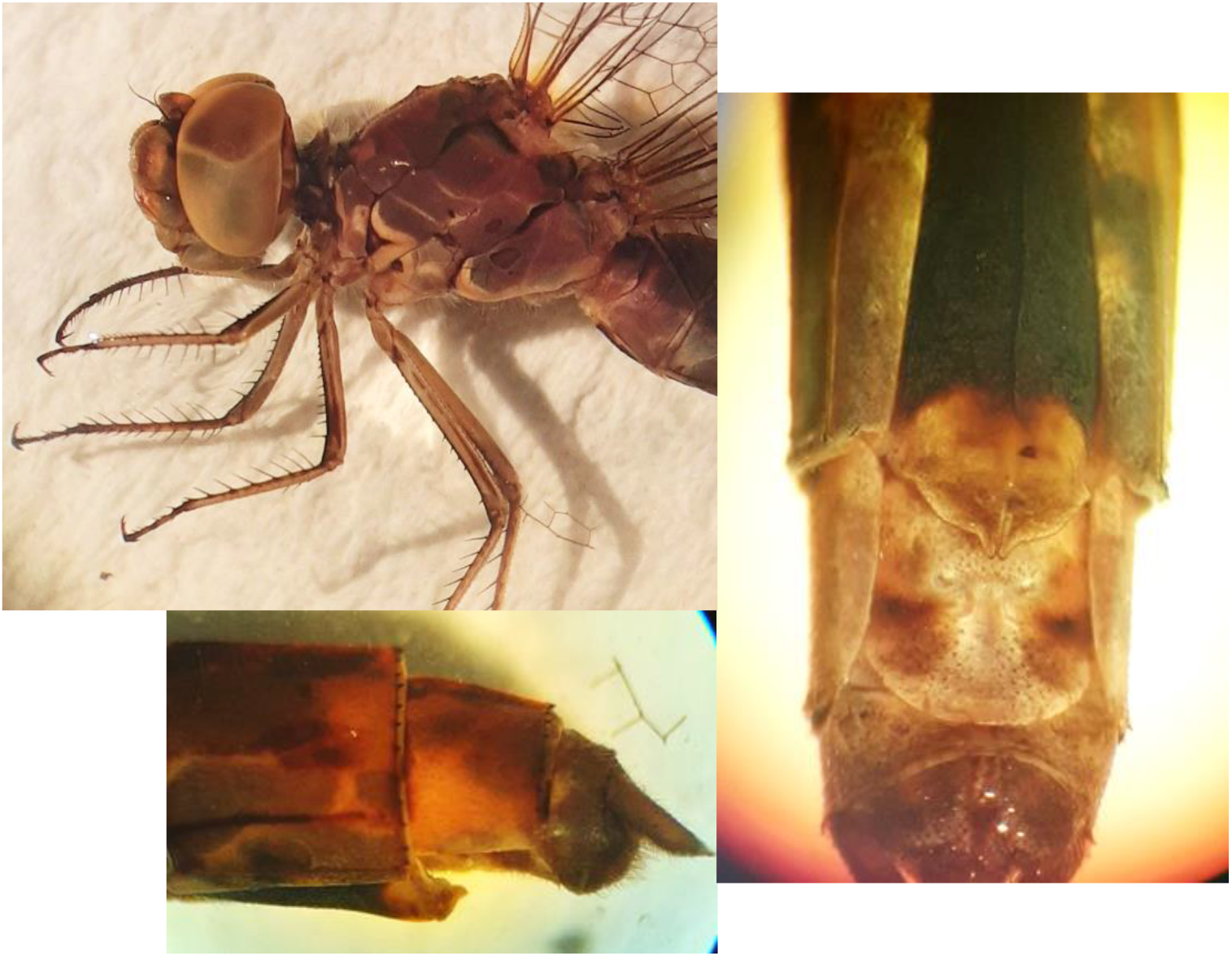
*Sympetrum meridionale* female, the second specimen from Cilieni Pool Băilești General habitus, vulvar scales in lateral and ventral view

4) *Sympetrum sanguineum* (Müller, 1764) vs. *Sympetrum bigeminus* (bioRxiv preprint 10.1101/2022.09.17.5508356).

## Discussions

The general habitus and the morphological characters of the collected specimens of *S. fonscolombii* and *S. striolatum* correspond to the descriptions in the odonatological literature (Askew 2004, Cîrdei & Bulimar 1965, Boudot et al. 2019, Dijkstra et al. 2020, Smallshire & Swash 2020, Wildermuth & Martens 2019).

For *S. sanguineum*, the vulvar scale is presented differently in the literature: not prominent, not bilobed / bifid (Askew 2004), pointed, not bifid Cîrdei & Bulimar (1965), pointed and bifid Dijkstra et al. (2020).

For *S. meridionale*, the vulvar scale is prominent (well visible) in lateral view and bifid in ventral view in both collected specimens. This aspect is different than in odonatological literature:

- “scarcely visible in lateral view” and illustrated straight, not bifid / bilobed (Askew 2004),
- “very small” and illustrated with a very small median point flanked by two reduced plates (Cîrdei & Bulimar 1965, fig. 222, pg. 246),
- Dijkstra et al. 2020 – vulvar scale has the same aspect as presented by Cîrdei & Bulimar (1965)

In one of the two females from Cilieni Pool (*S. meridionale*), the bifid vulvar scale is similar almost to identity with the vulvar scale of *S. sanguineum* ? *bigeminus* presented in the bioRxiv preprint 10.1101/2022.09.17.5508356 (Babalean 2022). The presence of an identical/very similar bifid vulvar scale in two females, one with black legs (typical for *S. sanguineum*) and the other with predominantly yellow legs (typical for *S. meridionale*) shows a problem. The genus Sympetrum has a problem at least in species diagnosability if not in species composition.

## Notes

### Competing Interest Statement

The authors have declared no competing interest.

